# SARS-CoV-2 Non-Structural Protein 1(NSP1) Mutation Virulence and Natural Selection: Evolutionary Trends in the Six Continents

**DOI:** 10.1101/2022.07.22.501212

**Authors:** Samira Salami Ghaleh, Karim Rahimian, Mohammadamin Mahmanzar, Bahar Mahdavi, Samaneh Tokhanbigli, Mahsa Mollapour Sisakht, Amin Farhadi, Mahsa Mousakhan Bakhtiari, Donna Lee Kuehu, Youping Deng

## Abstract

Severe acute respiratory syndrome coronavirus 2 (SARS-CoV-2) is an unsegmented positive-sense single-stranded RNA virus that belongs to the *β-coronavirus*. This virus was the cause of a novel severe acute respiratory syndrome in 2019 (COVID-19) that emerged in Wuhan, China at the early stage of the pandemic and rapidly spread around the world. Rapid transmission and reproduction of SARS-CoV-2 threaten worldwide health with a high mortality rate from the virus. According to the significant role of non-structural protein 1 (NSP1) in inhibiting host mRNA translation, this study focuses on the link between amino acid sequences of NSP1 and alterations of them spreading around the world. The SARS-CoV-2 NSP1 protein sequences were analyzed and FASTA files were processed by Python language programming libraries. Reference sequences compared with each NSP1 sample to identify every mutation and categorize them were based on continents and frequencies. NSP1 mutations rate divided into continents were different. Based on continental studies, E87D in global vision and also in Europe notably increased. The E87D mutation has significantly risen especially in the last months of the study as the first frequent mutation observed. The remarkable mutations, H110Y and R24C, have the second and third frequencies, respectively. Based on this mutational information, despite NSP1 being a conserved sequence occurrence, these mutations change the rate of flexibility and stability of the NSP1 protein, which can eventually affect inhibiting the host translation.

**IMPORTANCE:** In this study, we analyzed 6,510,947 sequences of non-structural protein 1 as a conserved region of SARS-CoV-2. According to the obtained results, 93.4819% of samples had no mutant regions on their amino acid sequences. Heat map data of mutational samples demonstrated high percentages of mutations that occurred in the region of 72 to 126 amino acids indicating a hot spot region of the protein. Increased rates of E87D, H110Y, and R24C mutations in the timeline of our study were reported as significant compared to available mutant samples. Analyzing the details of replacing amino acids in the most frequent E87D mutation reveals the role of this alteration in increasing molecule flexibility and destabilizing the structure of the protein.

## INTRODUCTION

Severe acute respiratory syndrome coronavirus 2 (SARS-CoV-2) is an unsegmented positive-sense single-stranded RNA virus that belongs to the *β-coronavirus* genus and *coronaviridae* family, first reported in Wuhan (1, 2). Phylogenetic analysis revealed that SARS-CoV-2 has a 79% sequence similarity with SARS-CoV that caused an outbreak in 2002 (3). Therefore, it is of vital importance to understand the differences and the relationships between these two types of viruses (4).

The SARS-CoV-2 genome is ~ 30 kb in length and composed of 13–15 (12 functional) open reading frames (ORFs) which are located 5’ to 3’ and known as ORF1a and ORF1b that encode 16 non-structural proteins (Fig. 1. B). In the following, several structural proteins such as spike proteins (S), envelope protein (E), membrane protein (M), and nucleocapsid protein (N) encode via the structural region of the genome (5, 6) that have a significant role in virus pathogenicity and infectivity (7). Among these structural regions, there are accessory factors such as ORF3a, ORF6, ORF7a, ORF7b, ORF8, and ORF10. This segmentation is displayed in Fig 1. A. These gene products play important roles in viral entry and RNA synthesis, fusion, and survival in host cells, and also have the capability of gaining rapid mutations as the virus spreads (8–10).

**FIG 1.**
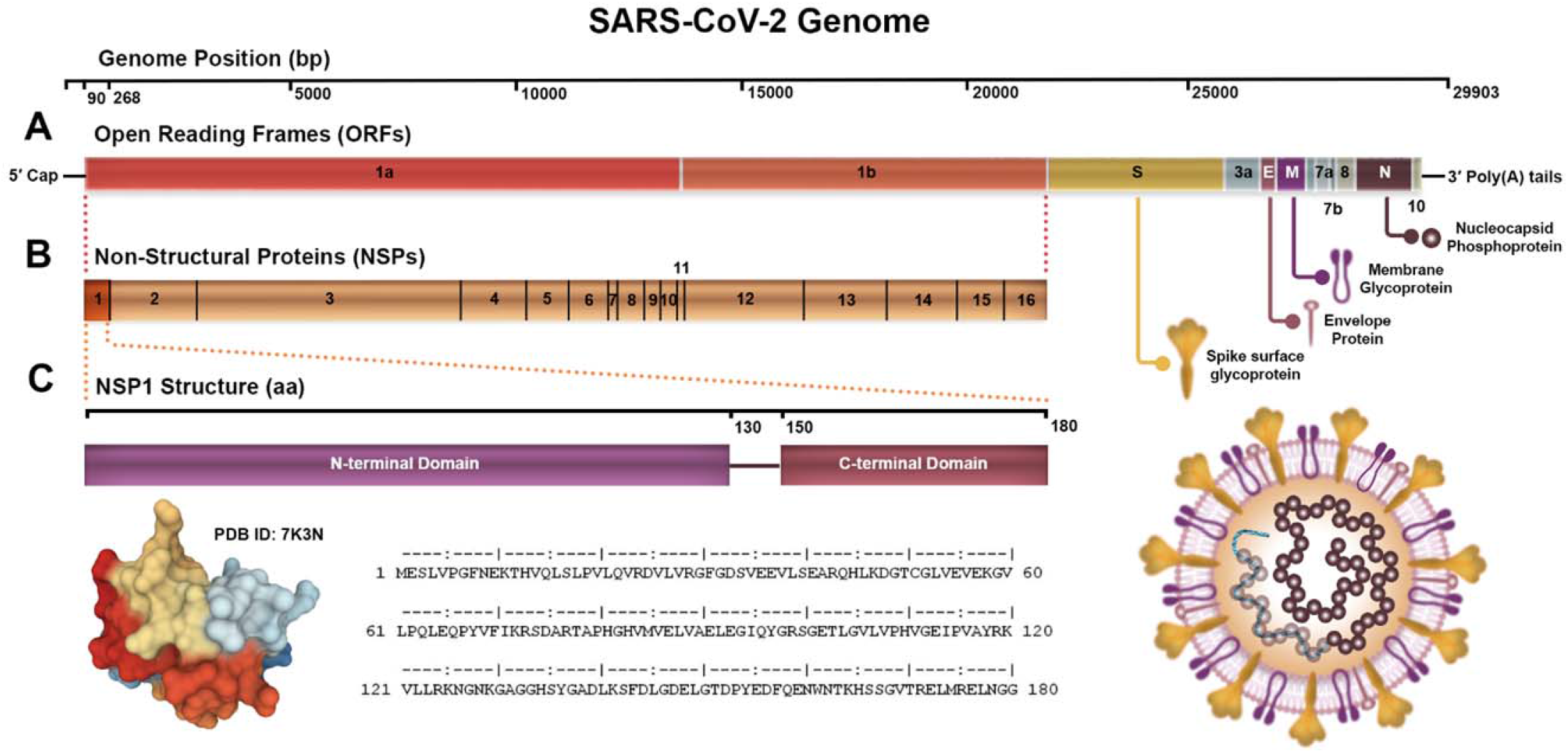
The schematic diagram illustrates an overview of the SARS-CoV-2 genome more detailed NSP1. (A) SARS-CoV-2 genome organization includes the ORF1ab sequence, the structural regions (S, E, M, and N) and accessory factors (ORF3a, ORF6, ORF7a, ORF7b, ORF8, and ORF10). (B) The ORF1ab sequence is self-cleaved into 16 non-structural proteins, and (C) NSP1 domains, 3D structure, and its amino acid sequence.

**FIG 2.**
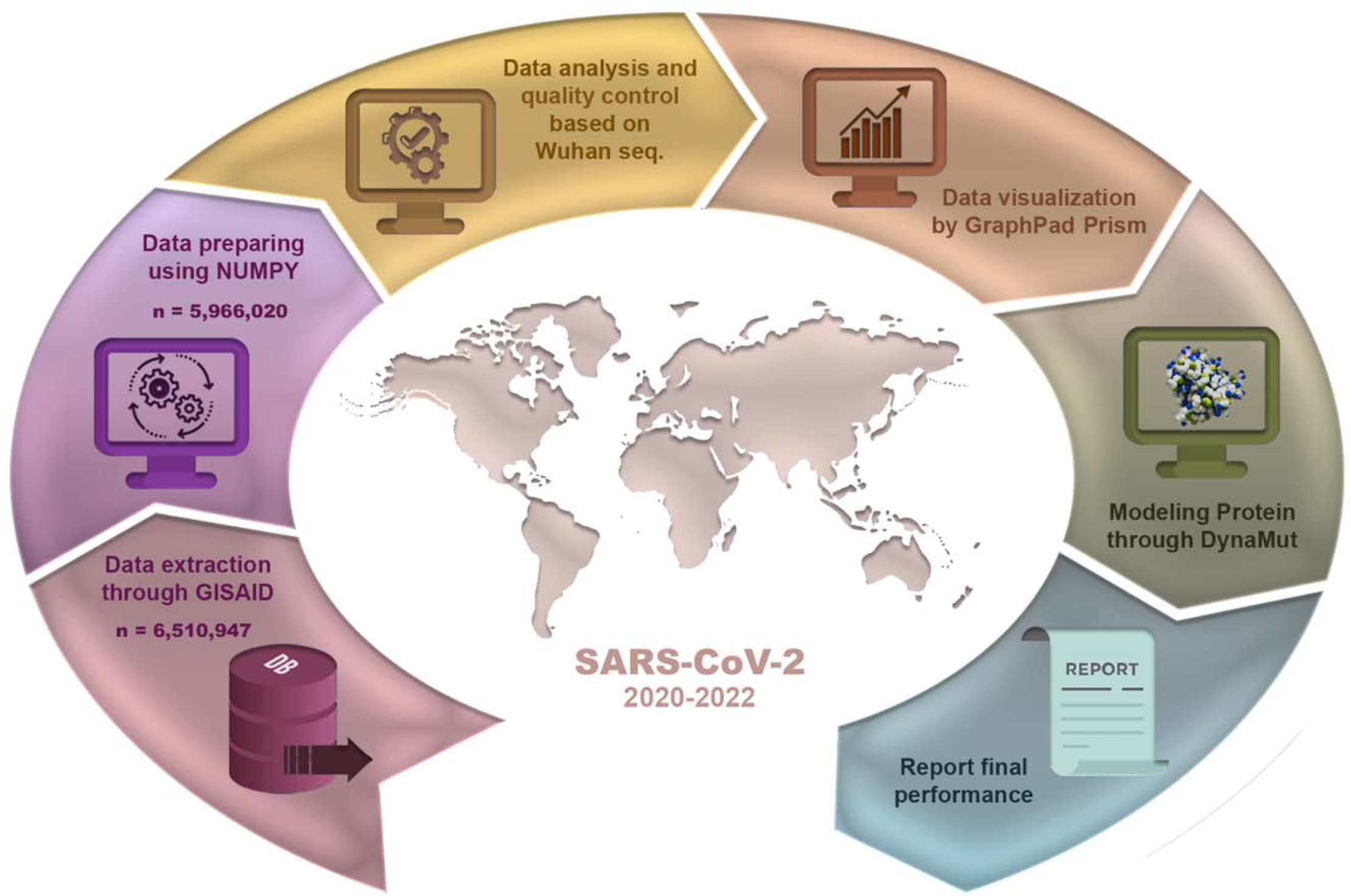
Data processing workflow steps are used to represent and validate NSP1 AAs mutations.

A major virulence factor of SARS-CoVs is the non-structural protein 1 (NSP1), which has a significant role in suppressing host gene expression by ribosome association. Binding NSP1 from SARS-CoV-2 to the 40S ribosomal subunit of the human cell as a host results in shutting down of mRNA translation both *in vitro* and *in vivo.* Cryo-electron microscopy analysis helps to clarify how they are placed together. When the NSP1 C-terminus (Fig. 1. C) binds to the 40S subunit of the ribosome (6), the formed complex prevents binding the mRNA to the ribosomal entry tunnel. NSP1 blocks gene I-dependent innate immune responses by retinoic acid-inducible that finally leads to eradication of the infection. The structural characterization of the inhibitory mechanism of NSP1 against SARS-CoV-2 can be useful for further significant insights into the drug design industry (11). NSP1 acts as a ribosome gatekeeper, binds the small ribosomal subunit and stalls canonical mRNA translation at various stages during initiation (12, 13). In addition, degradation of host mRNA occurs because of binding to the ribosome that leads to endonucleolytic cleavage (14). Thus, NSP1 is a major viral virulence factor and because of this sensitive role of NSP1 in Coronavirus disease 2019 (COVID-19) tracking any structural mutation is vitally important (15).

Over time, the increase in number of patients suffering from COVID-19 and mortality rate differed in geographical regions. This difference appears because of multiple reasons that can affect these studied mutations (16). Due to the modifications on NSP1 protein amino acids (AAs), the incidence of different variations on the SARS-CoV-2 was observed. These mutations can affect binding to the human 40S subunit in ribosomal complexes and then, change flexibility and stabilization in the virus. Translation inhibition takes place as a result of this phenomenon that occurs in both *in vitro* translation systems and *in vivo* (17).

Despite global vaccination efforts, various mutations continue to be observed that can represent virus escape from vaccination and all other administered drugs (eg. Remdesivir, hydroxychloroquine, chloroquine, ribavirin, ritonavir, lopinavir, favipiravir, interferons, bevacizumab, azithromycin, etc) (18). Location-dependent species in each continent demonstrate vaccination has not been able to have the expected effect of controlling the disease (19). Thus, having clear information from amino acid sequences (AASs) and the virus structure is useful and necessary to detect more details about the mutation dynamics on the SARS-CoV-2 genome (20).

In this regard, we aimed to investigate the amino acid mutation patterns and their specificities divided into geographical areas from the beginning of the pandemic to January 2022. In addition, we represented the link between mutations and incidence of them on different continents that can be helpful in the prediction of dynamic transmission in the future.

## RESULTS

### Numbers and incidence of mutations in NSP1 AAS based on geographical areas

Relevant statistical analyses were performed to identify the potential mutations in the NSP1 protein structure. At the present, the number of AAS mutations in the whole of 6,510,947 sequences was examined. Of the total number of NSP1 FASTA files, 277,242 data belong to not-matched length samples, 3,434 data relate to non-human sequences, and 264,251 data belong to sequences that contained X. Finally, 544,927 data were not included in the analysis, and 5,966,020 NSP1 data remained for examination in the study.

The statistical results showed that 6.3106% of AASs had one mutation, 0.2024% of AASs included two mutations, 0.0039% of AASs comprised three mutations, and 0.0010% of AASs contained more than four mutations. In contrast, 93.4819% of sequences had no mutation in their AASs. In addition, the frequency percentage of mutations was investigated in six geographical regions; North America, South America, Europe, Asia, Oceania, and Africa (Fig. 3. A and Supplementary file1).

**FIG 3.**
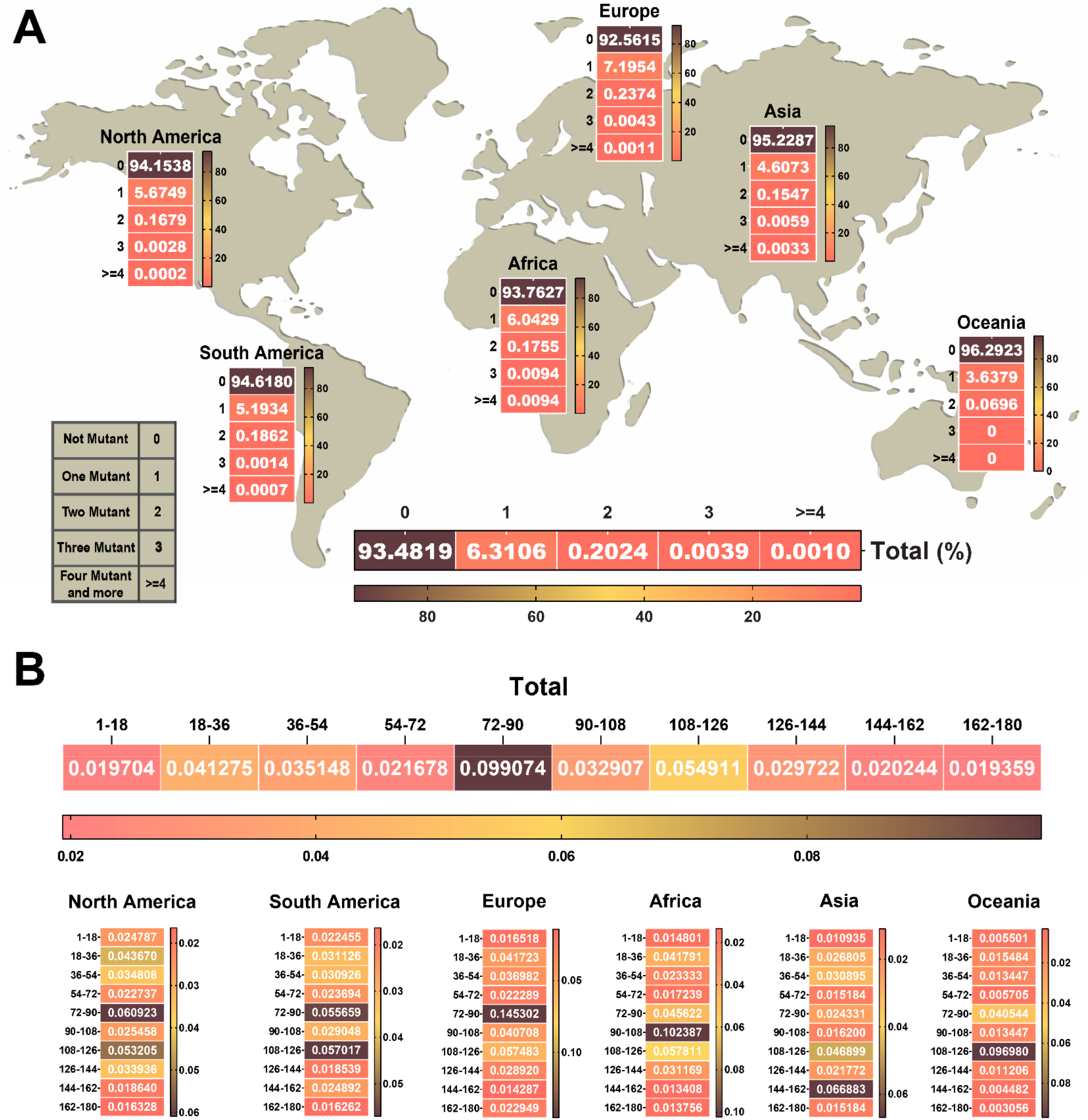
The heat map of the number and approximate region of mutations in NSP1 of SARS-CoV-2. (A) Heat map chart of the mutations’ number in NSP1 of SARS-CoV-2 until January 2022 in each continent. (B) The NSP1 protein was divided into ten segments. The rate of each mutation per segmented 18 amino acids was indicated in the heat map of the protein. The highest frequency rate occurred in the 72 to 126 amino acid sequence of the NSP1 protein that we can consider as a hot spot region in NSP1. The percentage of data were used to draw heat maps. Supplementary file1 and Supplementary file 2.

In continental-related analysis regarding the frequency of mutations number, Oceania has the least frequency of samples that have one or more than one mutations in their AASs. Also, the top rank frequency of no mutant samples is dedicated to this area. In this way, the data regarding Oceania with 27,270 AASs comprising of no mutations in 96.2923% of sequences, one mutation in 3.6379%, and two mutations in 0.0696% of AASs, but in the examined AASs from this continent, more than two mutations were not detected. Europe was labeled the top rank frequency of one and two mutations in samples compared to other regions. Despite that, Europe has the least frequency of no mutant samples. Data related to Europe indicates 2,931,894 AASs represent that no mutation was detected in 92.5615% of sequences, although one mutation in 7.1954% and two mutations in 0.2374% of their AASs were detected.

In Africa, we observed a high frequency of three or more than three mutations in samples. This continent with 31,907 AASs data revealed three mutations in 0.0094% of AASs, and more than four mutations in 0.0094% of AASs represented. The details about these frequencies are represented in Fig. 3. A.

According to the high percentage of no mutant regions on NSP1 AASs in all geographical areas, the data demonstrate NSP1 as a conserved region that was preserved against multiple mutagenic agents during the evolution of the virus. The mutation frequency was determined based on the number of mutations in each segment relative to the total mass. The worldwide results depicted that the highest mutation frequency occurred in the region of 72 to 90 AA (0.0990%) and then in the districts of 108 to 126 AA (0.0549%). More details about the frequency of mutations in each geographical area and the location where they belong separately were mentioned in Fig. 3. B and Supplementary file2.

### Mutation’s specificities according to geographical areas

More detail to survey the NSP1 AASs mutations, particularly the location of mutations in the protein structure and their frequency were investigated between January 2020 to January 2022. The first five frequent mutations regardless of geographical distribution were represented in Table 1.

**TABLE 1.** NSP1 top five frequent mutations from January 2020 to January 2022.

| Rank | Residue | Frequency (%) |
| --- | --- | --- |
| Top 1 | E(87)D | 1.1754 |
| Top 2 | H(110)Y | 0.3100 |
| Top 3 | R(24)C | 0.2937 |
| Top 4 | S(100)N | 0.1148 |
| Top 5 | E(37)D | 0.1104 |

The statistical incidence of these five mutations based on the continents is listed in Table 2. The significant point is all of these mutations didn’t appear among continents as the top mutations.

**TABLE 2.** The incidence of the global NSP1 top five mutations is based on the continents.

| Residue | Variant frequency (%) |  |  |  |  |  |
| --- | --- | --- | --- | --- | --- | --- |
|  | North America | South America | Europe | Asia | Oceania | Africa |
| E(87)D | 0.3475 | 0.1021 | 2.0835 | 0.1243 | 0.0806 | 0.1441 |
| H(110)Y | 0.3051 | 0.3862 | 0.3222 | 0.1946 | 1.5219 | 0.5077 |
| R(24)C | 0.3421 | 0.2229 | 0.2628 | 0.2001 | 0.1796 | 0.1723 |
| S(100)N | 0 | 0 | 0.1982 | 0 | 0 | 0 |
| E(37)D | 0.0728 | 0.1344 | 0.1480 | 0 | 0.0147 | 0.0752 |

Due to the results of our analysis, E87D in NSP1 was labeled the most frequent mutation up until January 2022 with a frequency rate of E87D at 1.1754%. Among these mutations, the E87D mutation was present on all continents except South America, as one of the top mutations. Therefore, Europe by 2.0835% of frequency is the highest, and Oceania by 0.0806% of frequency is the least ranked for E87D mutation. H110Y mutation has been observed as one of the top mutations with the highest frequency in Oceania and the least frequency in Asia by 1.5219% and 0.1946% of frequency respectively. R24C mutation has been remarked in North America as the highest prevalent mutation and in Africa as the least frequent mutation among the top mutations with 0.3421% and 0.1723% of frequency respectively. S100N mutation was detected only in Europe by 0.1982% frequency among the top mutations. E37D mutation has been shown in North America, South America, and Europe by 0.0728%, 0.1344%, and 0.1480% of frequency respectively.

In North America, R24C, E87D, and H110Y are the top three mutations that have more frequency than others, respectively. Also, there is more variety in the type of amino acid to which it is converted in the last ranks of mutation. South America has a different pattern in the frequency of the high-rank mutations. In South America, H110Y was labeled the first frequent mutation.

Regarding the Europe data, E87D is the first rank among mutations and there is a significant difference between the frequency of E87D and other mutations that have less frequency. This pattern has the most matches in the global ranking. In Asia, E148G is the most frequent mutation by 0.8849% of frequency that has a considerable discrepancy with other mutations. Indeed in the 148^th^ position of AAs, glutamic acid (E) replaces Glycine (G). Oceania especially has a remarkable role in enhancing the H110Y mutation global rate by 1.5219% of frequency that should be considered. In Africa, E102Q is the first frequent mutation. In this mutation, glutamic acid converts to glutamine with 0.9120% of frequency, and also lysine replaces glutamic acid with 0.5391% of frequency causing structural modification in AAs.

The global distribution in mutation rates represents that the NSP1 region is one of the conserved regions in the virus genome during evolution. In diverse geographical distributions, different percentages of mutations were observed that may lead to significant impacts on mortality rate, drug resistance or vaccine escape, and severity of the disease (21, 22). The top five mutations of each geographical area and which amino acids they have been replaced are shown in Fig. 4 based on the logarithm 10 of data frequency in the percentage of substituted AAs. The details about other happened mutations are available in the Supplementary file3.

**FIG 4.**
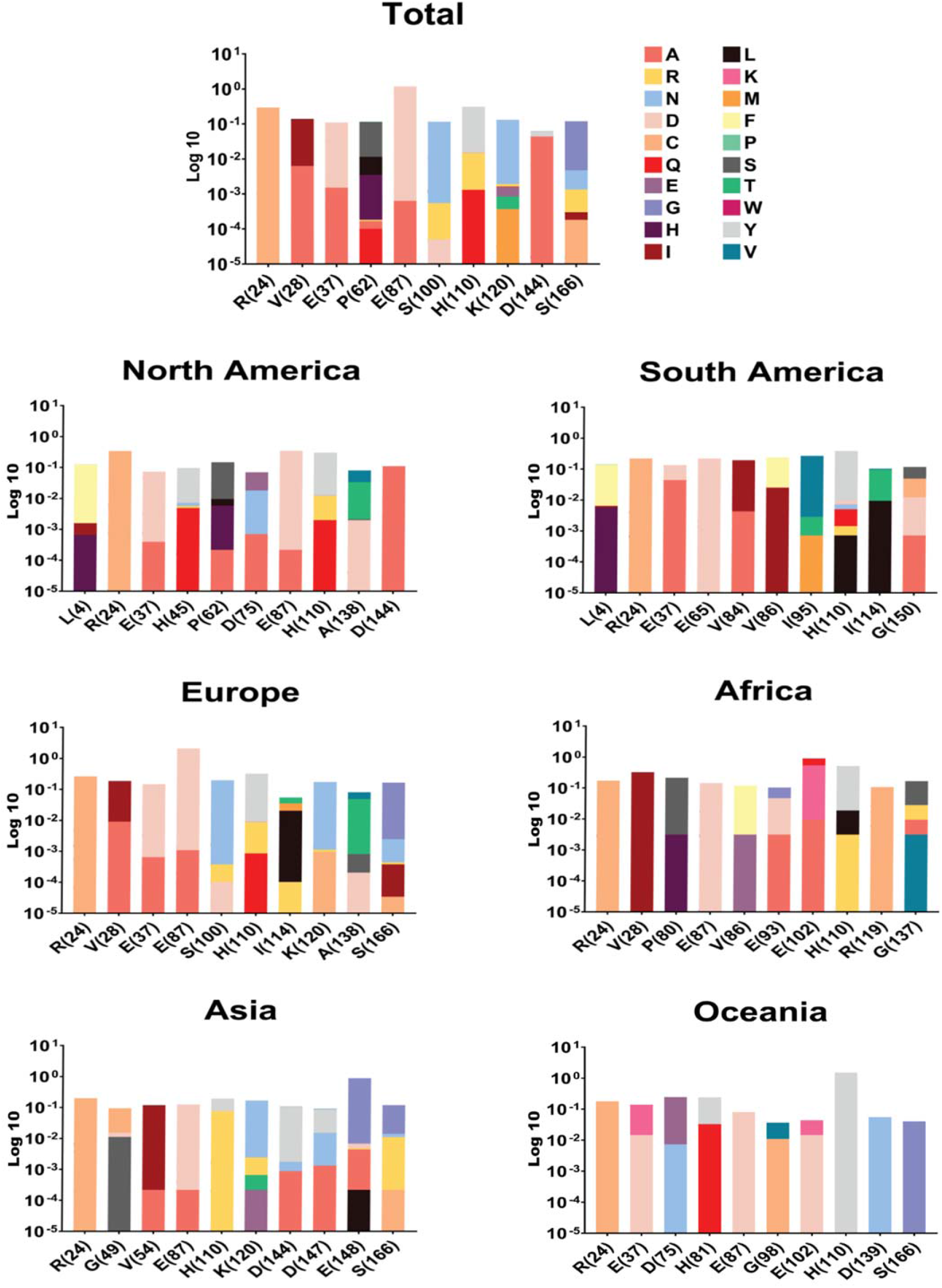
NSP1 top ten mutations with the highest frequency globally and geographic areas including North America, South America, Europe, Asia, Oceania, and Africa. The position of changed amino acids and substituted ones is shown differently based on the logarithm 10 of data frequency in the percentage of substituted AAs. The mutation frequency was evaluated for each of them by normalizing the number of genomes carrying a given mutation in an intended geographic area.

### Evolutionary assessment of happened mutations in the position of AAs of top five mutations according to time and geographical regions

To detect the incidence of each mutation, AASs of each geographic area were analyzed over time by classifying them according to the month of sample collection from January 2020 until January 2022, as represented in the GISAID database (23–25).

The continuation of mutations was investigated and collected monthly. In the following, fluctuations with a prevalence rate higher than 0.01 per AASs were reported. In the global timeline, two fluctuations of mutations that were incident in the R24 and E87 positions are more significant than the prevalence of other mutations. R24 had increased at the beginning of the pandemic which had a higher rate than other mutations and also had a peak in May 2020 to almost July 2020. On the other hand, the peak of E87 mutation started in May 2021.

Three high slope peaks of D75 (February 2020 by the highest rate 0.03), and R24 (May 2020, by the highest rate 0.02) have occurred in North America. D144 mutation peak incidence in this area is different from others and it was prevalent with less slope than the start in March 2020 and raised in September 2020 to 0.02 frequency. In South America, I114 has three continuous peaks between July 2020 and February 2021. V86 peaked in April 2021 up to 0.01 rates. In the last month of study in this area, we detected raising the frequency of V84.

In Europe, particularly E87 was increased and raised to a 0.03 frequency rate in July 2021. But in the last month of the study, the frequency increased. The timeline of Asia illustrated a peak of E148 in October 2021 with 0.09 frequency. Oceania has a peak of H110 mutation in May 2021 with the highest rate of 0.2. E102, the top mutation in Africa, reached a rate of 0.18 in November 2020. Detailed distribution of the five ranks in high mutation rates of NSP1 variants globally and on each continent is provided by the month of sample collection and displayed in Fig. 5. The details about other peaks of mutations that happened during the study timeline are also available in Supplementary file4.

**FIG 5.**
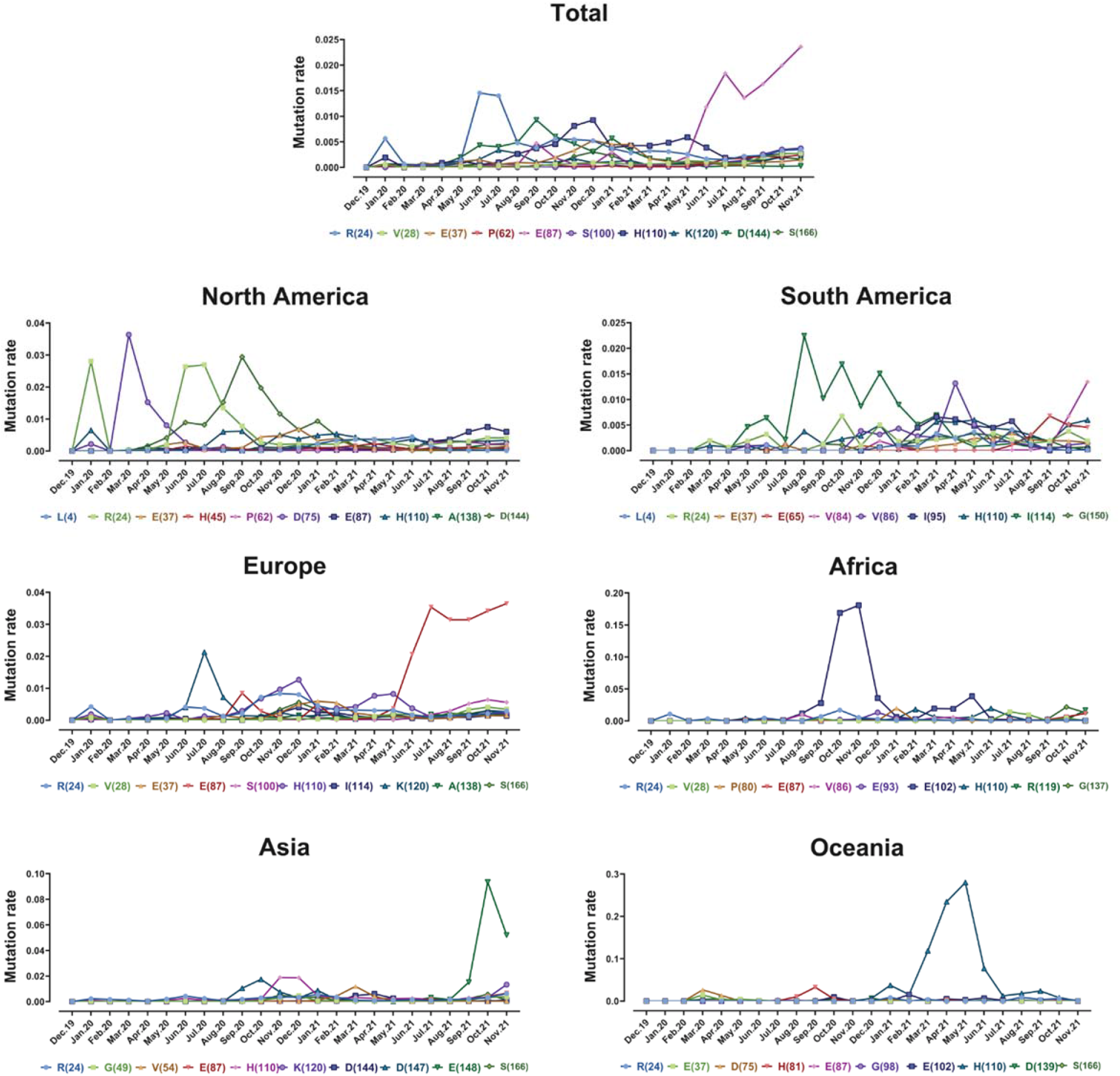
Plots time evolution pathways of top five high-rate mutations of NSP1 globally and in different geographic areas including North America, South America, Europe, Asia, Oceania, and Africa. The data was computed as the number of AASs having a given mutation over the total number of AASs according to the month of sample collection.

### Assessment of E87D mutation on dynamicity and flexibility of NSP1

To unveil the effect of E87D mutation on the structure of NSP1 protein, we used the DynaMut website for protein modeling. The variation in vibrational entropy energy (ΔΔS_vib_ENCoM) between wild and mutant types was calculated. The obtained data represent that the mutation at E87D increases the molecule flexibility on the NSP1 protein structure by a value 0.569 kcal.mol^-1^.K^-1^. On the other hand, binding affinity change caused by the alteration between glutamate and aspartate in position 87 can result in destabilizing the structure of NSP1 protein by a value −0.624 kcal/mol.

Investigation on the effects of this alteration in amino acids intramolecular interactions can give us a vision of the reason for destabilization in the protein structure after mutation.

Although both glutamate and aspartate are brønsted base and have a negative net charge in mutant type, by deleting a CH_2_ in the structure of residue, the three hydrogen bonds that made the protein more stable in wild-type were eliminated (Fig. 6).

**FIG 6.**
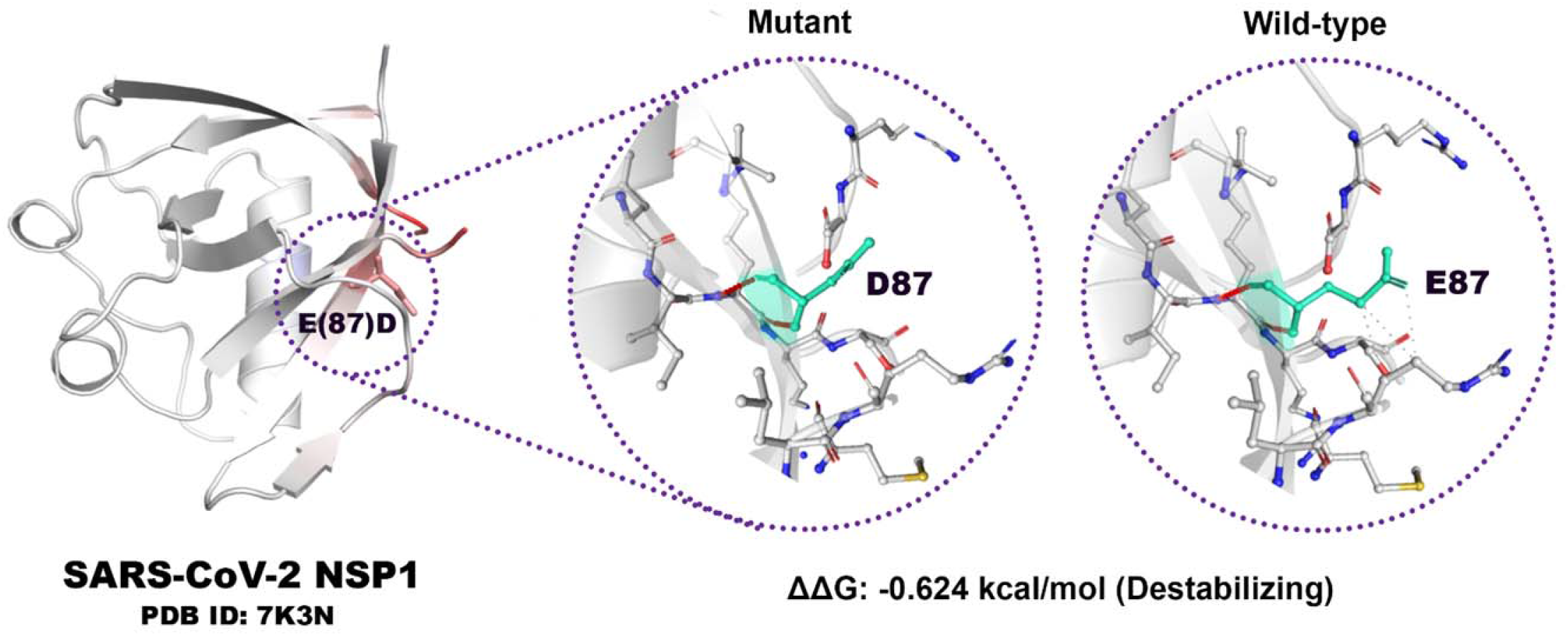
Impact of E87D mutation on structural dynamics of the NSP1 protein. Amino acids are colored according to the vibrational entropy change upon mutation. The red color represents a gain in flexibility of NSP1 protein. Wild-type and mutant residues are colored in light-green and are also represented as sticks alongside the surrounding residues which are involved in any type of interaction.

## DISCUSSION

SARS-CoV-2, as RNA viruses are more susceptible to mutations than DNA viruses(26). NSP1 is known as a critical virulent factor with significant effects on the virus-host interaction interface, such as inhibiting host mRNA translation (27, 28), antagonizing interferon (IFN) signaling (29), and inducing inflammatory cytokines and chemokines (30).

Furthermore, some of these effects on NSP1 are conserved in other β-CoVs and even α-CoVs. Given the very likely structural similarity between SARS-CoV-2 NSP1 and SARS-CoV NSP1, the information of SARS-CoV NSP1 functions could be highly helpful for understanding the biological and pathological roles of NSP1 in SARS-CoV-2 (31).

This study was designed to analyze all mutations that occurred between January 2020 to January 2022, and regional patterns of mutations on NSP1 detected in the following six regions; North and South America, Europe, Asia, Oceania, and Africa. A recent investigation demonstrated the effects of tirilazad, phthalocyanine, and zk-806450, which showed lower energy scores compared to alisporivir and cyclosporine, two compounds with *in vitro* activity against NSP1 that may have higher inhibition effectiveness (32). Hence, it is expected inhibiting NSP1 activity in the virus by these compounds leads to resuming host translation normally.

Esculin is a glucoside and naturally occurs in barley, and horse chestnut. It is given to improve capillary permeability and fragility and has been reported to inhibit collagenase and hyaluronidase enzymes. The molecule has been shown to have antioxidant and antiinflammatory activity (33). Esculin interacts mainly with R62, S63, A68, H72, and M74 (major interacting residues in the docking pose) through H-bond interactions with esculin. According to the preformed analysis, these interactions are located in the hotspot region of NSP1 that cause the influence of esculin on NSP1 protein function. This suggests the ability of esculin to not only inhibit NSP1 activity but also play a role against secondary symptoms such as inflammation. Hence, designing a drug against NSP1 in addition to other methods for treatment of COVID-19 likely gives a hopeful insight into possible control and finally eradicating the disease (34).

As for previous studies, E87D mutation is located on the globular domain of *in vitro* protein that can affect the stability and flexibility of the virus. Therefore, this mutation can help in gaining flexibility and destabilization in the protein structure. Based on a recent study, a decrease in molecular flexibility was detected in H110Y NSP1 as the second frequent mutation and D75E which is one of the less frequent mutants. When the R24C, as the third frequent mutation, is happening in NSP1, flexibility was changed in the region containing AA residues E65, L64, Q63, P62, C24, D16, and V14. Similarly, in some less frequent mutants like D48G NSP1, an increase in molecular flexibility was detected at AA residues S40, R43, Q44, K47, and G48. These alerts in AAS might have effects on its natural function in terms of blocking host mRNA translation and evading the immune system (35).

Different reasons involve natural selection in determining the fate of mutations in each continent (36). Depending on conditions and every factor that affects the viral genome, different mutations with different symptoms in patients are possible. Mutations that have enough potential of resistance against the host immune system, can adapt to the conditions and then, increase proliferation, and facilitate virus transmission.

Based on our results, there are some less frequent mutations such as S100N, E37D, V28I, K120N, and P62S in NSP1. But due to evolutionary factors, based on our results, there are some less frequent mutations such as S100N, E37D, V28I, K120N, and P62S in NSP1. However, there is no significant increase in the frequency of these mutations, while the frequency of obvious mutation E87D, notably has increased. The top five mutations position on AASs of NSP1 protein were shown in Fig. 7.

**FIG 7.**
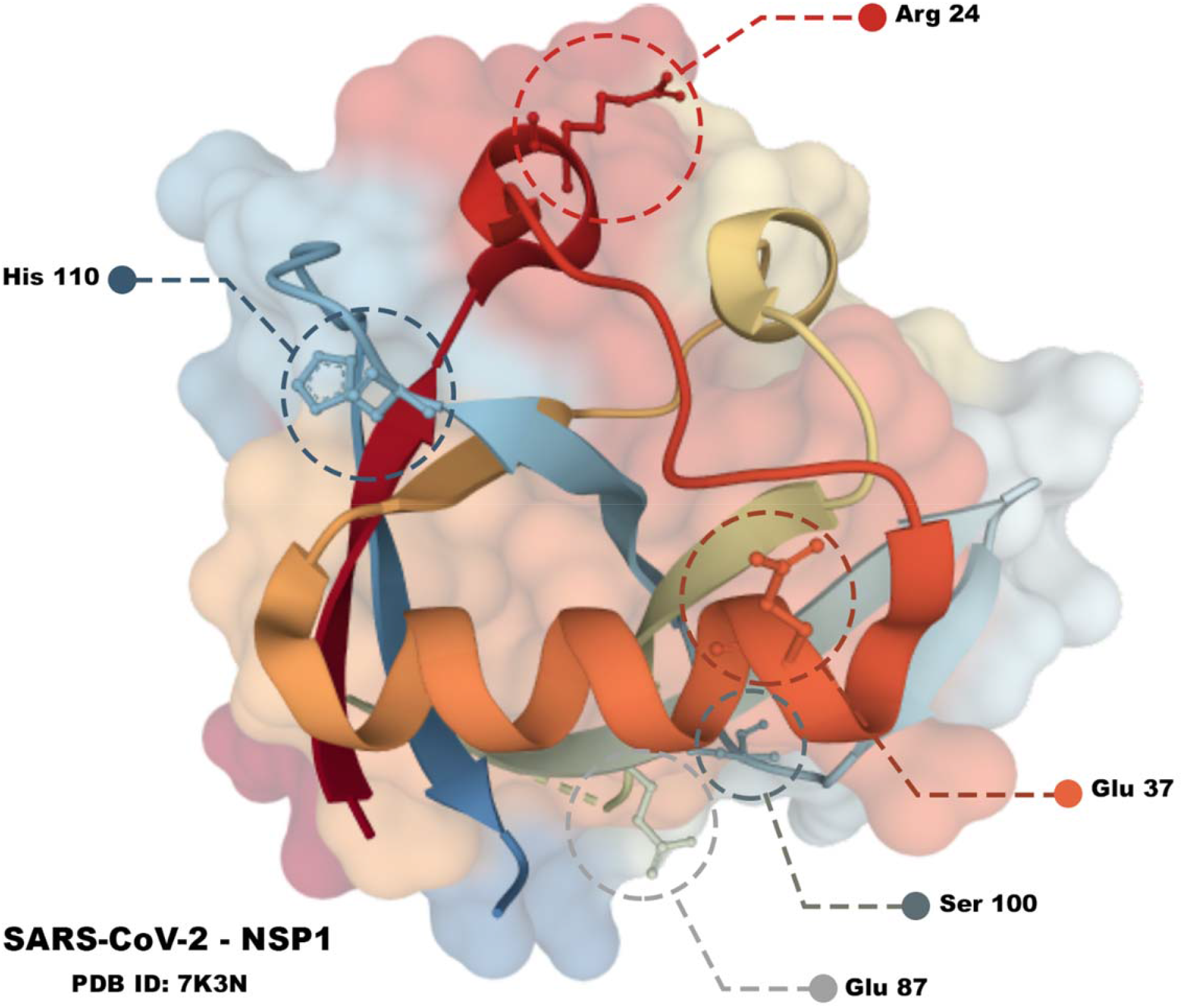
3D structure of NSP1 protein. The positions of top five frequent mutation on the protein structure are displayed in different colors (E87, R24, H110, S100, E37).

The tissue and cell expression patterns of known SARS-CoV-2 interacting human proteins, based on transcriptomics and antibody-based proteomics in the Human Protein Atlas database (https://www.proteinatlas.org/humanproteome/sars-cov-2) helped us to find the human genes that are related to NSP1 of SARS-CoV-2. COLGALT1, PKP2, POLA1, POLA2, PRIM1, and PRIM2 are the genes that encode collagen beta(1-O)galactosyltransferase 1, plakophilin 2, the catalytic subunit of DNA polymerase alpha 1, the accessory subunit of DNA polymerase alpha 2, DNA primase subunit 1, and 2 respectively. The products of these genes have low immune cell specificity and commonly medium tissue expression. Generally, these genes play a role in DNA replication except COLGALT1 and PKP2. Collagen beta(1-O)galactosyltransferase 1 by acting on collagen glycosylation facilitates the formation of collagen triple helix and plakophilin 2, which may play a role in junctional plaques (37–42).

There are two limitations in the present study that should be noted. One of the limitations is focusing only on AASs without considering the nucleotide sequences. Another limitation in this process is that samples reported from Europe had more rates than the data which were reported from other regions, and reciprocally Oceania is the one area that has the least sample rate. Thus, there may be other significant mutations present without extensive sequencing of samples that have not been available in the data.

Our findings suggest mutations of SARS-CoV-2 are expanding and help the virus adapt inside of hosts, providing conditions for virus survival and making other mutations in new geographical areas. E87D, H110Y, and R24C mutations increased in the timeline of the study as the first, second, and third frequent mutant sequences, respectively. The effects of these mutations on the flexibility and stabilizing of NSP1 and subsequently on survival features of the virus can have different results in patients that should be considered. To figure out the precise impact of the mutations on the disease, further studies should be done in the future.

## MATERIALS AND METHODS

### Sequence and Source

This study focuses on evaluating a big database of NSP1 AASs of the SARS-CoV-2 genome. NSP1 is located on the N-terminal region of the ORF1ab sequence that is susceptible to many mutations. The reference sequence is the Wuhan-2019 virus with access number ‘EPI_ISL_402124’ also known as wild type. All of the AAS samples were compared with the reference sequence. 180 AAs of the NSP1 region were extracted from GISAID (www.gisaid.org) (23–25) database belong period January 2020 to January 2022. Erasmus Medical Center grunted the accession to this database.

### Sequence analyses and processing

NSP1 was extracted from other genes and after analyzing mutations and sequence alignment, FASTA files were processed by Python 3.8.0 software. ‘Numpy’ and ‘Pandas’ libraries optimized the whole process. Mutations were defined as each difference between sample and reference, then the location and replaced AA were reported. Less or more than 180 AAs belong to non-human samples such as pangolin or bat, and samples containing nonspecified AAs (reported as X) were eliminated, and finally, 6,510,947 refined samples were examined in the study. The identifying algorithm for detecting mutants is as follows:

In this algorithm, since all sequences have equal lengths, respectively ‘Refseq,’ and ‘seq’ refer to reference sequence and sample sequence.

for refitem, seqitem in zip (refseq, seq)
if (refitem! = seqitem)
Report a new mutant

Each sample’s continent name and geographical coordinates were labeled on them. Pycountry-convert 0.5.8 software and ‘Titlecase’ library in Python were used to report global prevalence maps of mutations. The procedure of data refining was presented in Fig. 2.

### Secondary protein structure and dynamic prediction

Analyzing the mutational structure and molecular flexibility of NSP1 protein modeling was performed by the DynaMut web server (http://biosig.unimelb.edu.au/dynamut/). The PDB ID of protein (7K3N) was taken from the Protein Data Bank (https://www.rcsb.org/) and used for prediction on the DynaMut web server.

### Statistical analysis

R 4.0.3 and Microsoft Power BI software were used to conduct data normalization and comparison charts outlining. GraphPad Prism 8.0.2 software were used for data visualization. To improve and compare the data’s results, the normalized frequency of each region was reported. Then the number of mutations was divided by the number of sequences on that region comparable in equal proportions.

## List Of Abbreviations

(SARS-CoV-2): Severe acute respiratory syndrome coronavirus 2
(ORF): Open reading frames
(S): Spike proteins
(E): Envelope protein
(M): Membrane protein
(N): Nucleocapsid protein
(NSP1): Non-structural protein 1
(COVID-19): Coronavirus disease 2019
(AAS): Amino acid sequence
(AA): Amino acid
(IFN): Interferon

## Supplementary

- Supplementary file1.xlsx
- Supplementary file2.xlsx
- Supplementary file3.xlsx
- Supplementary file4.xlsx

## Acknowledgments

The authors thank all of the researchers who have shared genome data openly via the Global Initiative on Sharing All Influenza Data (GISAID).

## Authors’ contributions

K.R., M.M.S., M.M.B., and A.F. contributed to data collection. Y.D., M.M., and K.R. contribute to study design, K.R., and M.M. design workflow and code, and data analysis. B.M, K.R., and M.M., and A.F. contributed to data visualization. S.S.G. wrote the manuscript. S.T., M.M., D.L.K. and B.M. corrected the manuscript and provided useful comments. M.M., K.R., and B.M. monitored the accuracy of Additional data. B.M. designed graphical contents. Y.D. have final edited and supervised the work.

## Data availability

The raw data supporting the conclusions of this article is available in supplementary file(s).

